# Transient active osmotic swelling of epithelium upon curvature induction

**DOI:** 10.1101/2020.11.25.398107

**Authors:** Caterina Tomba, Valeriy Luchnikov, Luca Barberi, Carles Blanch-Mercader, Aurélien Roux

## Abstract

Generation of tissue curvature is essential to morphogenesis. However, how cells adapt to changing curvature is still unknown because tools to dynamically control curvature in vitro are lacking. Here we developed self-rolling substrates to study how flat epithelial cell monolayers adapt to a rapid, anisotropic change of curvature. We show that the primary response is an active and transient osmotic swelling of cells. This cell volume increase is not observed on inducible wrinkled substrates, where concave and convex regions alternate each other over short distances, identifying swelling as a collective response to changes of curvature with persistent sign over large distances. It is triggered by a drop in membrane tension and actin depolymerization, perceived by cells as a hypertonic shock. Osmotic swelling restores tension while actin reorganizes, probably to comply with curvature. Epithelia are thus unique materials that transiently, actively swell while adapting to large curvature induction.

---

Many morphogenic events are characterized by epithelium folding: gastrulation^1,2^, neurulation^3^ and villification of the intestine^4^ are driven by cell shape changes caused by changes of cell surface tension such as apical constriction^5–7^. Cell surface tension results from plasma membrane tension, acto-myosin contractility and cell-cell and cell-substrate adhesion forces^8^, and is thus a major regulator of morphogenesis.

Folding can also occur through compressive forces within epithelia^9^, forming buckled shapes predicted by mechanical models^10^ and produced by several processes: proliferation under externally imposed confinement^11^, differences of proliferation between domains of the same tissue^12,13^, or active contractility of muscles around a growing intestinal epithelium^14^. Thus, mechanical stresses emerging in reaction to external constraints are determinant of epithelium folding.

However, very little is still known about how tissues respond in time to external mechanical stresses. Cell monolayers under compression maintain their cell density constant through time by increasing cell extrusion and reducing cell proliferation^15^. Opposite reactions are seen upon tissue stretching^16,17^. These results concern flat tissues and do not address how cells react to changes of the 3-dimensional (3D) tissue organization, in particular to changes of their curvature.

Curved substrates have been developed for cell culture^18^ in order to study curvotaxis^19^, a term that designates all processes guided by curvature such as cell shape, motility, adhesion and gene expression. For instance, concave surfaces orient cell polarity and cell migration^20,21^, increasing migration speed by reducing contacts with the substrate^22^. In contrast, convex regions enhance osteogenic differentiation by modulating cytoskeleton forces acting on the nucleus^22^. At a multi-cellular scale, curvature controls acto-myosin contractility during tissue growth^23^, epithelium detachment from concave regions^24,25^, and collective migration inside and outside of cylindrical substrates^26,27^. All these curvature and actin dependent cell processes require to integrate local cell constraints over long distances through cell-cell and cell-substrate adhesions^28^. However, all these studies investigate cells adhering on substrates with fixed curvature, while *in vivo*, they experience forces and curvatures that are constantly changing.

In order to study how tissues would react to induction of curvature, we looked for techniques that could acutely change the curvature of the substrate onto which cells would have been grown. Engineering of controlled strain gradients in nanomembranes leading to self-rolling of thin solid films^29,30^ offered a versatile strategy to produce complex 3D structures for a large panel of applications in flexible electronics, soft robots and biomedical devices^31,32^. This approach was recently used in biomimetic self-folding macro-structures, based on anisotropic swelling of printed hydrogel composites^33^. Biocompatible micro-containers were developed to control drug release^34,35^, to encapsulate cells in rolled films for potential implantable tissue grafts^36^, or to mimic blood vessels and muscle fibers^37^. These bioengineering technologies^38^ proved cell viability on scaffolds that spontaneously change their shape^39^. Here, we present a self-rolling elastomer thin film that forces a cell monolayer to fold into a tubular shape.

We prepared self-rolling substrates according to the fabrication process reported in Fig. 1a. The technique is based on built-in spontaneous curvature of a polydimethylsiloxane (PDMS) film^40^, presents the advantage of creating anisotropic curvature, and provides an internal control, as the PDMS film does not roll over parts strongly adhering to the bottom substrate. It consists of a superposition (from bottom to top) of a PDMS layer on a glass slide, a gelatin film circumscribed to the central part of the PDMS layer below, and finally a bilayer of two different PDMS composites: the top leaflet of PDMS contains silicon oil and is strongly adherent to the bottom, oil-free PDMS leaflet. Oil extraction is performed through immersion of the whole substrate in isopropanol, forming a pre-constrained PDMS bilayer, which spontaneously rolls when cut over the gelatin^40^. The PDMS bilayer adheres to the bottom PDMS outside the gelatin patch, keeping it flat until cut.

**Figure 1.**
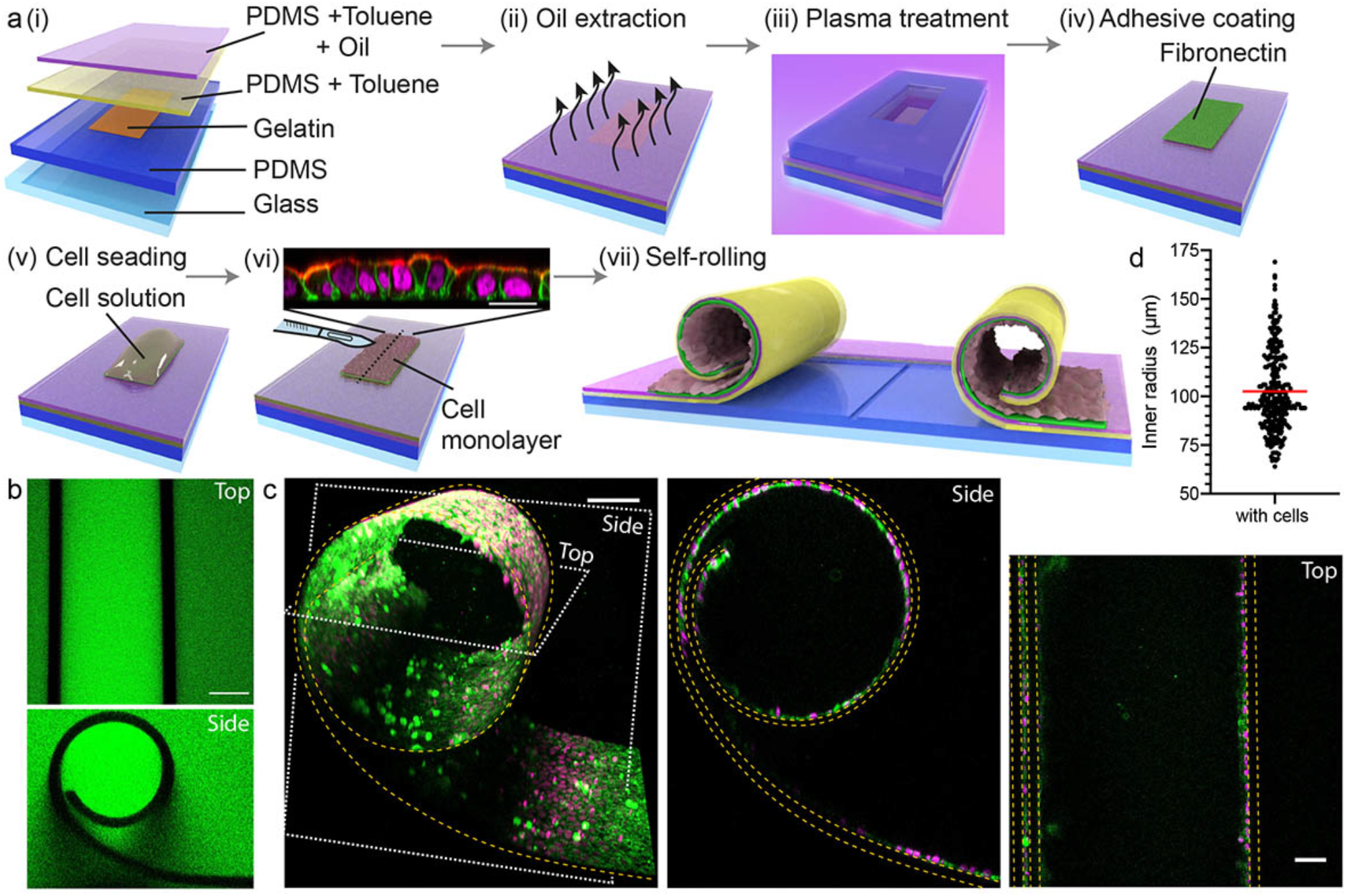
Self-rolling substrates induce cylindrical deformation of epithelial cell monolayers. **a**, Schematic of concave curvature generation: (i) multi-layer substrate composed of: glass slide (light blue); PDMS sheet (dark blue); central gelatin Patch (brown); bottom PDMS leaflet and toluene solution (yellow); top leaflet of PDMS, toluene and silicon oil solution (pink); (ii) oil extraction by isopropanol bath and formation of the strain gradient; (iii) surface activation by plasma treatment on the central region through a PDMS mask; (iv) fibronectin solution (green) incubation on the central region; (v) cell solution (light pink) incubation; (vi) substrate cutting after cell monolayer formation (inset: orthogonal view of a polarized cell layer, ezrin (red), cell membrane (Myr-Palm-GFP, green), and nuclei (H2B-mCherry, magenta). Scale bar=20 μm); (vii) self-rolling of the substrate upon cutting. **b**, Top (top) and side (bottom) views of a PDMS tube without cells, dextran solution (green). Scale bar=50 μm. **c**, 3D view (left, scale bar=100 μm), side (middle) and top (right) views (scale bar=50 μm) of a PDMS tube with cells, cell membrane (deep-red CellMask, green), and nuclei (H2B-mCherry, magenta). **d**, Distribution of inner radii of tubes with cells. The red line stands for the mean value, n=278 images; N of independent replicates=3.

To confine cell adhesion, a delimited area is activated by plasma surface treatment, and then coated with fibronectin. We employed Madin-Darby Canine Kidney (MDCK) cells that form well-organized epithelial layers, which cell mechanics and collective behavior is among the best described. Cells were let to adhere and form a polarized monolayer for 24h. Finally, by cutting the PDMS bilayer, and consequently the cell monolayer adhering to its top surface, the pre-constrained PDMS bilayer is allowed to roll (Fig. 1a, Supplementary Movie 1).

Multi-rolls were discarded because cells may be squeezed between two (or more) roll layers, adding an undesired constraint (Supplementary Fig. S1). We thus optimized the formation of single roll tubes (Fig. 1b,c) by reducing the region covered by gelatin (down to 0.5cm width) and limiting the cut length (up to 1.5cm). As the radius of curvature is proportional to the substrate thickness^40^, we fixed a PDMS bilayer thickness of 14±1 μm (Supplementary Fig. S1) that produces relevant curvature radii (Fig. 1d) found *in vivo*. We observed that the tube radii were maintained constant over time (Supplementary Fig. S1), suggesting a constant constraint on the cell monolayer. Using a simple elastic model (Supplementary Information, Section 1), we estimated that the substrate exerts a pressure of ~4*Pa* onto the monolayer while imposing its preferred radius of curvature. The cell monolayer is not expected to significantly unroll the PDMS substrate, given that it is much less rigid than the PDMS. In particular, we predicted that the relative deviation of the tube radii from the optimal radius of curvature of the substrate due to the elastic resistance of the cell monolayer is in the order of 4% (Supplementary Information, Section 1).

Moreover, we estimated the change of thickness of the PDMS bilayer and of the epithelial cells upon rolling (Supplementary Information, Section 2). If purely incompressible (constant volume), cells are expected to thicken by 25% and decrease their width by 20%. If cells are compressible, their volume should decrease by ~5% (Supplementary Information, Section 1). We then investigated these predictions.

We first analyzed changes of cell shape upon rolling. To combine high-statistics, a time resolution of few minutes after rolling and a high-resolution imaging, cells were chemically fixed with 4% paraformaldehyde (PFA) at different timepoints after rolling. We used MDCK stably expressing the nucleus marker H2B-RFP and the plasma membrane marker Myr-Lyn-GFP^41^ or MDCK labelled with the plasma membrane marker CellMask and the nuclear stain Hoechst (see Methods). To analyze their shape, we segmented cells using Limeseg, a 3D cell segmentation ImageJ plugin^42^, and we quantified the average cell volume over time, both on planar and tubular regions (Fig. 2a). Cell volumes and shapes on flat PDMS do not change after fixation, showing that PFA does not affect our analysis (Supplementary Fig. S2).

**Figure 2.**
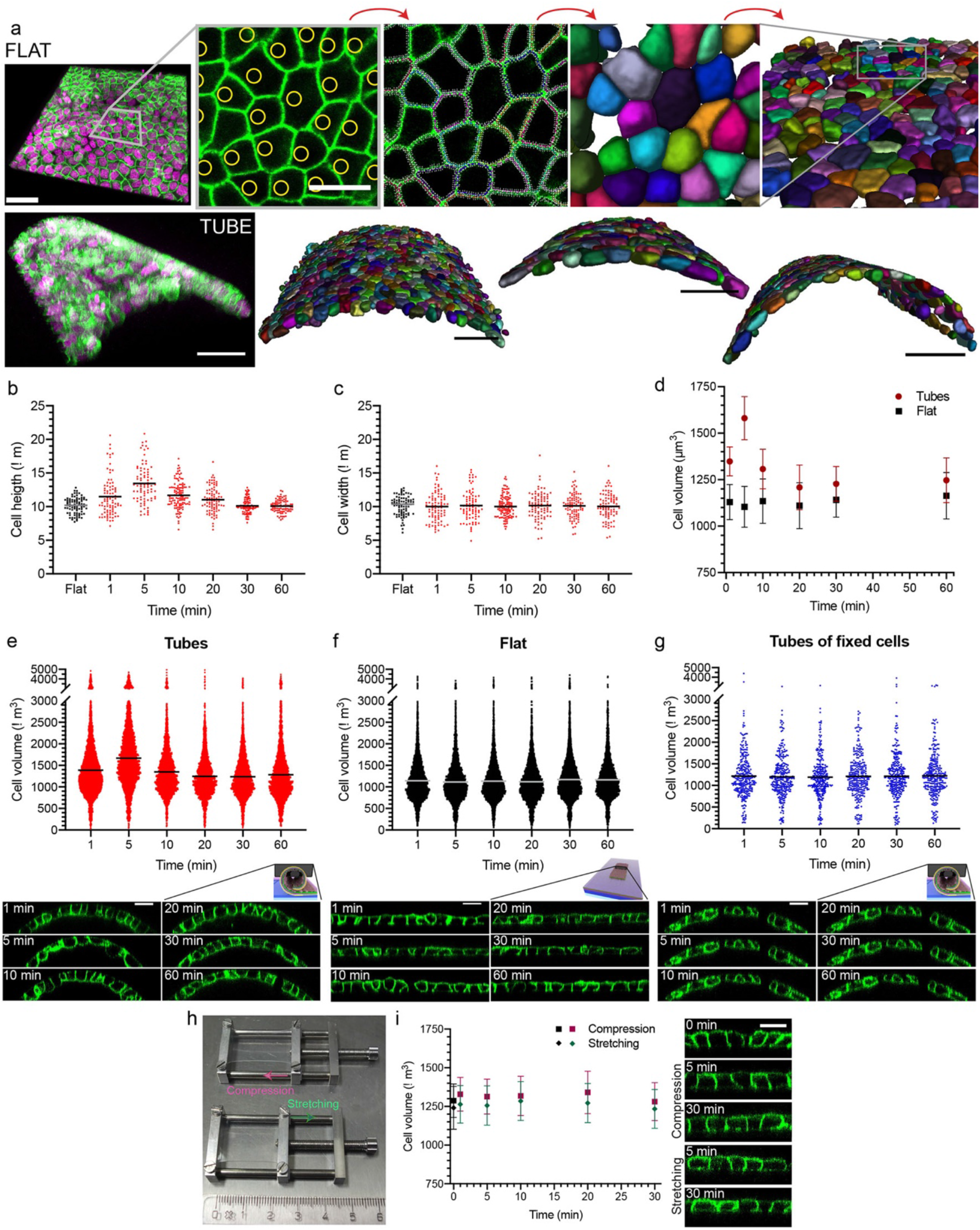
Cell volume increase and recovery are an active response. **a**, Cell volume measurements. Top, from left to right: 3D view of a cell monolayer on a flat PDMS substrate (scale bar=50 μm), cell membrane (Myr-Palm-GFP, green), and nuclei (H2B-mCherry, magenta); centroid identification of the nuclei (yellow circles) by Imaris segmentation (scale bar=20 μm); 3D cell membrane identification by LimeSeg segmentation; 3D reconstruction of cell shapes (zoom and large views, random colors represent different cells). Bottom, from left to right: 3D view of a cell monolayer on the top region of a PDMS tube; 3 representative examples of 3D reconstructions of cell volume on PDMS tubes. Scale bars=50 μm. **b**, Distribution of cell height on flat regions (black squares) over time (minutes) and on tubes (red circles) over time after cutting. n≥81cells/timepoint; N≥3. **c**, Distribution of cell width on flat regions (black squares) over time and on tubes (red squares) over time after cutting. n≥81cells/timepoint; N≥3. **d**, Weighted means and standard error of the weighted mean (SEWM, with variance weights) of cell volume on flat regions (black squares) over time and on tubes (red circles) over time after cutting. n≥13images/timepoints; N≥4. **e**, Distribution of cell volume on tubes (red circles) over time after cutting. n≥2700cells/timepoint; N≥4. **f**, Distribution of cell volume on flat regions (black circles) over time. n≥3200cells/timepoint; N≥4. **g**, Distribution of cell volume on tubes formed after cell fixation in PFA (blue circles) over time after cutting. n≥279cells/timepoint; N=1. Insets in **e-g**: representative orthogonal views of cells on tubes or flat regions (see diagrams) at different time points after cutting, cell membrane (Myr-Palm-GFP, green). Scale bars=20μm. **h**, Pictures of the stretchers in the initial position before compression (top) and stretching (bottom) of about 15% of the elastomer film. 1 small mark in the ruler = 1 mm. **i**, Weighted means and SEWM (with variance weights) of cell volume on an elastomer film before compression (black square) or stretching (black rhombus) and after compression (magenta squares) or stretching (green rhombuses) over time. n≥14images/timepoint; N≥3. N is the number of independent replicates, and the horizontal lines stand for the mean values.

Orthogonal views of the cell monolayer showed that cell thickness increased and peaked at 5min after rolling (Fig. 2b), while the cell width stayed constant (Fig. 2c). This cell shape change led to a mean aspect ratio (height/width) of 1.40±0.47 at 5min, compared to 1.03±0.23 on flat regions (Supplementary Fig. S2). The resulting cell thickening is thus higher and delayed in time compared to the maximum value of 25% predicted for a rolled incompressible epithelium (Supplementary Information, Section 2). Moreover, the cell apical pole, normally very flat (Fig. 2f), appeared swollen (Fig. 2e). This was associated with a volume increase up to ~40% at 5min (Fig. 2d,e), compared to cell volume on flat regions (Fig. 2d,f). This volume increase is fully consistent with the observed ~40% increase of height without width reduction (Fig. 2b,c). This is an unusual response of material to bending constraints, observed in rare materials with negative compressibility such as some rear crystals^43^, polymer foams^44^ and bi-material ligaments^45^.

However, and different from materials with negative compressibility, the gradual increase was followed by a slow recovery to normal volume within 20min, suggesting that the response and its recovery were active (energy-consuming) cell processes. Other than volume increase, we observed no other striking change of cell shape. We observed neither cell displacements (Supplementary Movie 2), nor cell shape elongation in the axis of the tube (Supplementary Fig. S2), nor increased cell extrusion (Supplementary Movie 3).

To test if volume increase and its recovery were active responses to curvature, we first investigated the deformation of cells chemically fixed before rolling. No cell volume increase was observed in this case (Fig. 2g), confirming that epithelial cells actively increased their volume upon rolling. This result also confirmed that the volume increase observed after rolling in living cells was not resulting from the roll geometry altering quantification of cell volumes. We then wondered whether the response could be induced by compressive or stretching stresses occurring upon curvature induction. We estimated the theoretical compression of the PDMS layer upon rolling to be less than ~15% (Supplementary Information, Section 3). We used mechanical stretchers to produce cell compression and stretching while the overall shape of the epithelium stayed flat. To apply compression, cells were grown on a pre-stretched PDMS layer by ~15%, which was then released. Inversely, to produce stretching, cells grew on a PDMS layer that was then stretched by ~15% (Fig. 2h). In both conditions, cell volume stayed constant over time (Fig. 2i, Supplementary S2), supporting that cell volume increase was specific to curvature induction.

Finally, we confirmed that cell volume stayed constant after cutting a flat epithelium, implying that cell volume increase was not a consequence of damages to the cells (Supplementary Fig. S2). Altogether, our results show that the transient cell volume increase is an active response to rapid change of curvature.

We next wondered which parameters of curvature could directly impact the cell response. As shown in Fig. 1d, the mean curvature radius obtained in the tubes with a cell monolayer was 102.7±20.9μm. Interestingly, the peak of cell volume around 5min occurred independently of the curvature radii (Fig. 1d), as long as it was below a threshold radius of ~180μm (Supplementary Fig. S3). Density also affected the volume change, as for half seeding densities, cell volume remained constant (Supplementary Fig. S3), suggesting that volume increase depended on cell compaction.

Surface curvature has two possible orientations (convex vs. concave), and we wondered if both orientations could produce a similar volume increase. To roll the PDMS bilayer in the other orientation, we inverted the order of the two PDMS leaflets (Fig. 3a). With this approach, the curvature radii obtained were larger (Fig. 3b). However, we still observed a cell volume increase upon substrate rolling (Fig. 3c), associated with apical height increase (Fig. 3d), and a recovery within 20min. Thus, in rolls, whether the induced curvature is positive or negative triggers the same active response of the epithelium, i.e. a transient increase in cell volume.

**Figure 3.**
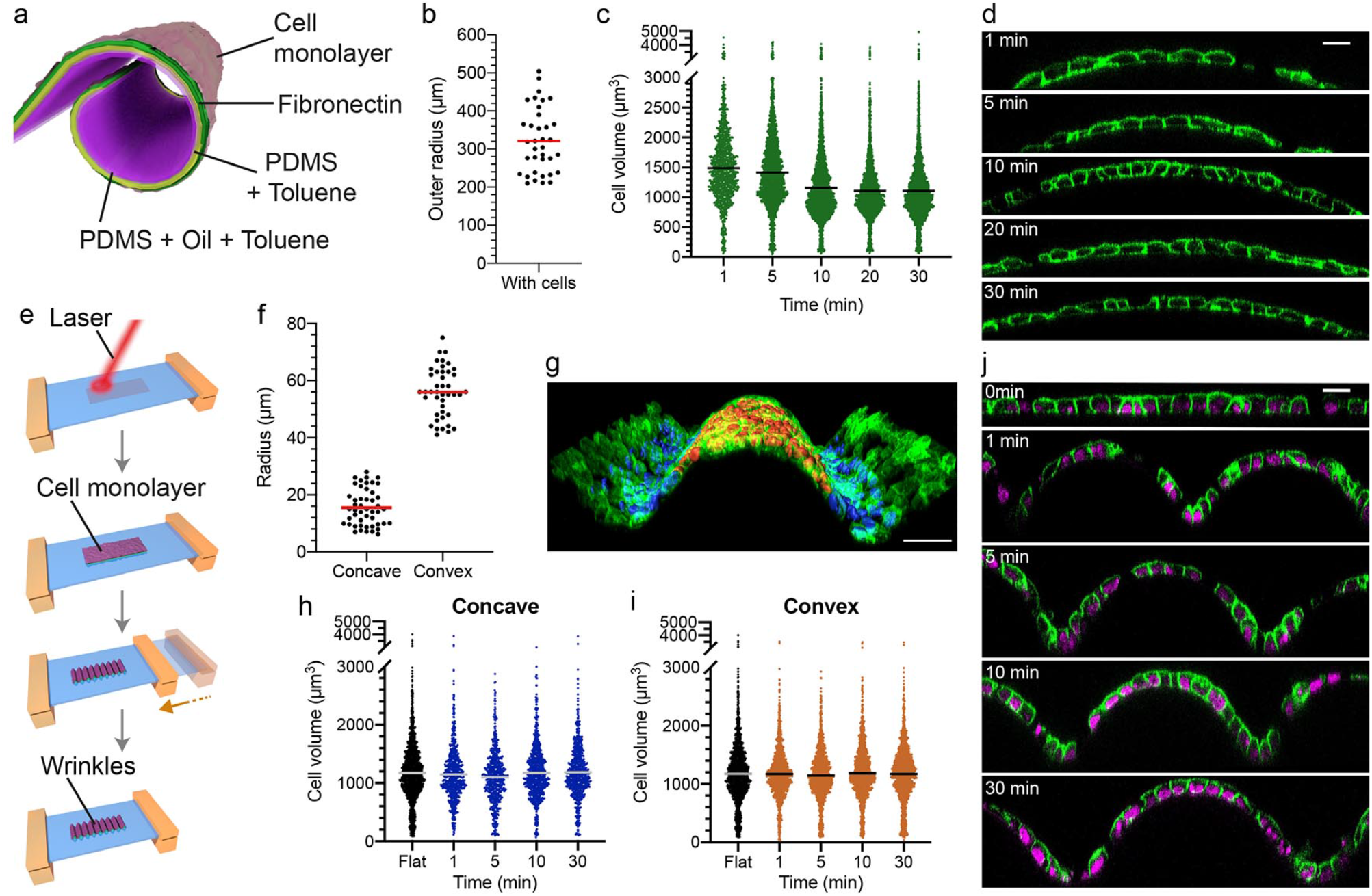
Both signs of curvature produce transient cell shape changes but this effect is compensated on wavy substrates. **a**, Schematic of convex curvature generation: PDMS and toluene with (pink) and without (yellow) silicon oil leaflets are inversed, compared to Fig. **1a**. Fibronectin coating (green) and cell monolayer (light pink). **b**, Distribution of outer radii of tubes with cells. n=36images; N=2. **c**, Distribution of cell volume on tubes with inverted sign of curvature (green circles) over time after cutting. n≥1179cells/timepoint; N=3. **d**, Representative orthogonal views of cells on tubes with inverted sign of curvature at different time points after cutting, cell membrane (Myr-Palm-GFP, green). Scale bar=20 μm. **e**, Schematic of wavy substrate formation. From left to right: infra-red laser exposure of the pre-stretched elastomer film; cell solution incubation on the flat surface of the pre-stretched elastomer film; wavy morphologies at the surface result from the stress relaxation of the substrate. **f**, Distribution of radii of the elastomer film in the concave and convex regions. n=48 (concave) and 43 (convex) images; N=1. **g**, 3D view of a representative cell monolayer on a wavy substrate, cell membrane (Myr-Palm-GFP, green), 3D reconstruction of nuclei segmented with Imaris on the concave (blue) and convex (orange) regions. Scale bar=50 μm. **h**, Distribution of cell volume on flat regions before substrate relaxation (black circles) and concave regions of wavy substrates (blue circles) over time after relaxation. n≥602cells/timepoint; N=1. **i**, Distribution of cell volume on flat regions before substrate relaxation (black circles) and convex regions of wavy substrates (orange circles) over time after relaxation. n≥1240cells/timepoint; N=1. **j**, Representative orthogonal views of cells on flat and wavy regions at different timepoints after relaxation, cell membrane (Myr-Palm-GFP, green), and nuclei (H2B-mCherry, magenta). Scale bar=20 μm. N is the number of independent replicates, and the horizontal lines stand for the mean values.

In rolls, curvature is the same over long distances. We thus wondered how the response of the epithelium would be when both curvatures are induced on the same substrate. To do this, we took advantage of a technique that we recently developed to produce wavy substrates^41^. First, a thin hard layer at the surface of a prestrained elastomer is formed by infrared laser irradiation, which then spontaneously adopts a wrinkled morphology upon stress relaxation (Fig. 3e). Radii of curvature obtained with this technique (Fig. 3f) are locally smaller than the values given by the self-rolling substrates (Fig. 1d, 3b). We first grew cells until confluence on flat pre-stretched and irradiated substrates, and relaxed the substrate to form the wrinkles. Then, cells were chemically fixed and segmented at different time points after wrinkling. We quantified cell volumes in the concave and convex regions (Fig. 3g), and observed that they did not change (Fig. 3h,i). Although cells in the concave regions looked highly deformed (Fig. 3j), these regions were only a few tens of microns, supporting the notion that the same curvature has to be propagated over long distances to have a significant impact on cell volume. Furthermore, in wrinkles, the effect induced by positive curvature could be compensated by the surrounding negative curvature. Overall, our results show that the cell volume increase is a collective response to induction of curvature over large spatial regions.

To find cellular functions involved in cell volume increase after rolling (Fig. 2), we looked at processes known to participate into cell volume regulation. We identified a number of ionic channels that are involved in volume regulation during osmotic shocks and beyond^46^, the cell cytoskeleton and TOR (Target Of Rapamycin) complexes 1&2 signaling, which are major players of cell volume and surface regulation^47^. Not surprisingly, both the cytoskeleton and TORC1&2 are also part of the cell volume response to osmotic shocks^48,49^.

We first inhibited volume-regulated anion channels^50^ (VRAC) by treating cells with DCPIB^51^. DCPIB-treated cells slowly increased their volume upon rolling, but without reaching the peak observed in non-treated cells (Fig. 2d). Only partial recovery was observed after 30min (Fig. 4a,b, Supplementary Fig. S4). Other key cell volume regulators are the Na^+^-H^+^ exchangers (NHE), whose activity was blocked by EIPA. Inhibition of NHEs, known to regulate volume increase^52^, led to a reduced volume increase upon rolling, which fully recovered after 30min (Fig. 4c,d, Supplementary Fig. S4). We then used Bumetanide to inhibit Na^+^-K^+^-2Cl^−^ cotransporter channels (NKCCs), which maintain the cell osmotic balance. Bumetanide treatment also caused a lower initial volume increase, but also strongly affected the volume recovery (Fig. 4e,f, Supplementary Fig. S4). Similar results were obtained with Furosemide, another common NKCC blocker (Supplementary Fig. S4). We concluded that ion channels were involved in both the initial volume increase after rolling, and its recovery. We thus hypothesized that the transient cell volume increase could be osmotic. To test this possibility, we first applied hypotonic shocks on flat cell monolayers and probed if cell volume and membrane tension changed. To follow membrane tension, we employed FliptR (Fluorescent lipid tension Reporter, or Flipper-TR), a probe that reports changes in membrane tension by changing its fluorescence lifetime^53^. After hypotonic shock, cell volume increased to reach a plateau about 15min after the shock (Fig. 4g). Fluorescence lifetime of FliptR – and thus tension – followed the same dynamics (Supplementary Fig. S4). Notably, the volume change was due to an increase in cell height without changes in cell width (Fig. 4h), as seen in rolls (Fig. 2b,c), yet with a lower increase (Fig. 2d). As expected, hyper-osmotic shocks on flat epithelia caused an instantaneous lifetime decrease (Supplementary Fig. S4), associated to a volume decrease of ~50% with no recovery (Fig. 4i, Supplementary Fig. S4). The global cell organization on flat monolayers did not change, showing that volume change was again only accommodated through changes of cell height (Fig. 4i). These results established that osmotic shocks could change cell volume and membrane tension in epithelia. To test directly if osmosis was involved in the transient cell thickening after rolling, we intended to counteract this response by applying a concomitant hyperosmotic shock: cells dramatically flattened (Fig. 4j), preventing cell volume increase observed in isotonic conditions. These results supported the hypothesis that curvature-induced cell volume increase was due to changes of the osmotic balance.

**Figure 4.**
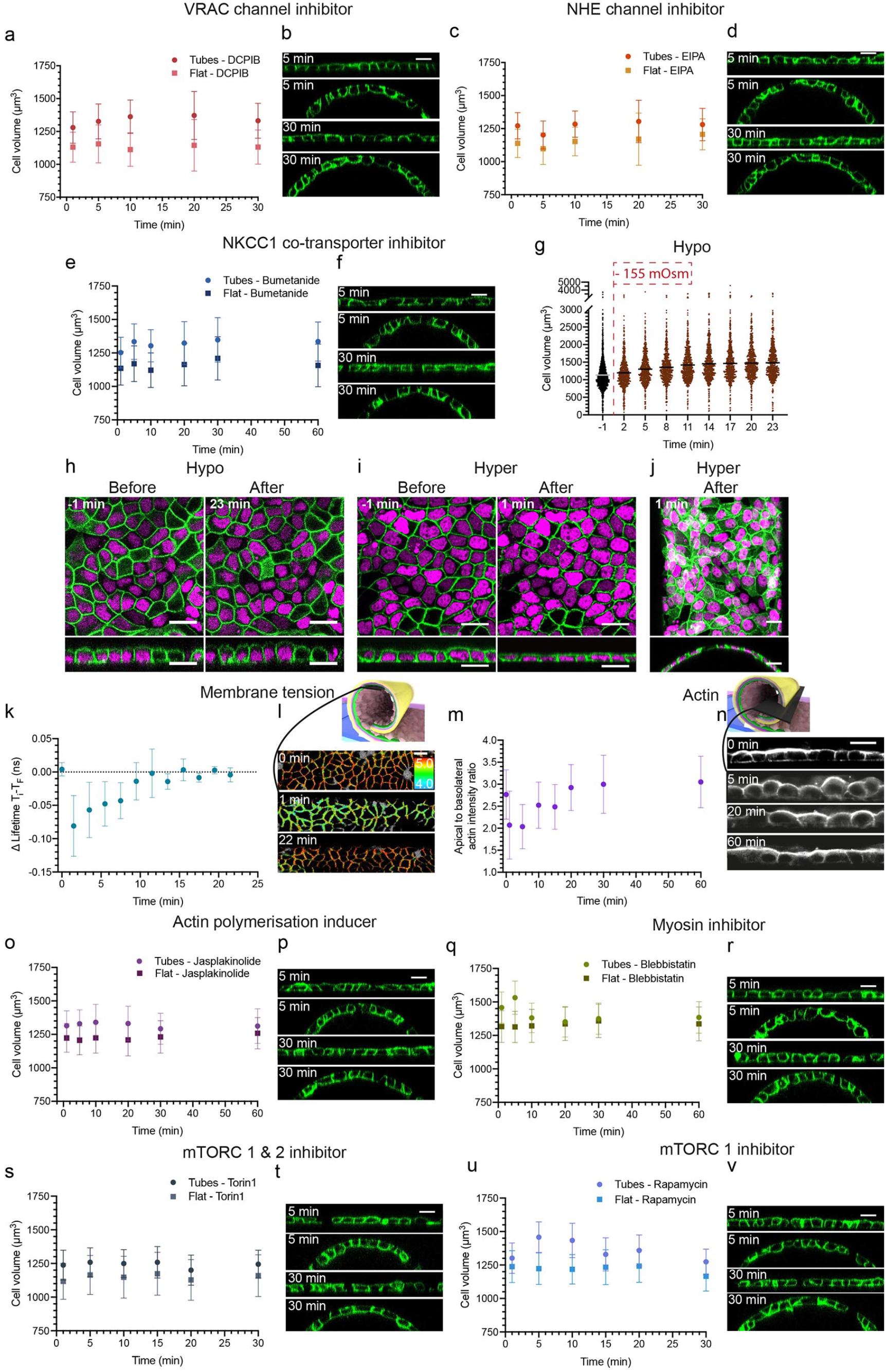
Ion channels, actin cytoskeleton and membrane tension control short and long-time scale response to curvature generation. **a**, Weighted means and SEWM (with variance weights) of cell volume after DCPIB treatment on flat regions (light pink squares) over time and on tubes (dark pink circles) over time after cutting. n≥6images/timepoint; N=3. **b**, Representative orthogonal views of DPIB-treated cells on flat regions or tubes at 5 and 30min after cutting. **c**, Weighted means and SEWM (with variance weights) of cell volume after EIPA treatment on flat regions (dark yellow squares) over time and on tubes (light orange circles) over time after cutting. n≥6images/timepoint; N=3. **d**, Representative orthogonal views of EIPA-treated cells on flat regions or tubes at 5 and 30min after cutting. **e**, Weighted means and SEWM (with variance weights) of cell volume after Bumetanide treatment on flat regions (dark blue squares) over time and on tubes (blue circles) over time after cutting. n≥9images/timepoint; N=3. **f**, Representative orthogonal views of Bumetanide-treated cells on flat regions or tubes at 5 and 30min after cutting. **g**, Distribution of cell volume on flat substrates before (black circles) and over time (minutes) after switching to a hypotonic solution (-~155 mOsm, brown circles). n≥596cells/timepoint; N=1, and the horizontal lines stand for the mean values. **h**, Representative images of cells on flat regions before and after hypoosmotic shock (-~155 mOsm), top (top) and side (bottom) views. **i**, Representative images of cells on flat regions before and after hyperosmotic shock (+ ~510 mOsm), top (top) and side (bottom) views. **j**, Representative images of cells on a tube after hyperosmotic shock (+ ~600 mOsm), maximum-intensity z-projection (top) and side (bottom) views. In **h-j**: Cell membrane (Myr-Palm-GFP, green), and nuclei (H2B-mCherry, magenta). **k**, Mean values and SD of fluorescence lifetime (difference values of each timepoint – T_i_ – with the final timepoint – T_F_) on flat regions (0min) and on tubes over time after cutting. n≥4images/timepoint, N=4. **l**, Representative top views along the tubes (see diagram) at different time points after cutting. The color bar corresponds to lifetime in nanoseconds (ns). **m**, Mean values and SD of apical to basolateral actin intensity ratio on flat regions (0min) and on tubes over time after cutting. n≥31images/timepoint; N≥3. **n**, Representative orthogonal views along the tubes (see diagram) at different time points after cutting. Actin (SiR-Actin, grey). **o**, Weighted means and SEWM (with variance weights) of cell volume after Jasplakinolide treatment on flat regions (dark violet squares) over time and on tubes (light violet circles) over time after cutting. n≥14/images/timepoint; N≥3. **p**, Representative orthogonal views of Jasplakinolide-treated cells on flat regions or tubes at 5 and 30min after cutting. **q**, Weighted means and SEWM (with variance weights) of cell volume after Blebbistatin treatment on flat regions (dark green squares) over time and on tubes (green circles) over time after cutting. n≥16images/timepoint; N≥3. **r**, Representative orthogonal views of Blebbistatin-treated cells on flat regions or tubes at 5 and 30min after cutting. **s**, Weighted means and SEWM (with variance weights) of cell volume after Torin1 treatment on flat regions (grey squares) over time and on tubes (dark grey circles) over time after cutting. n≥10images/timepoint; N=3. **t**, Representative orthogonal views of Torin1-treated cells on flat regions or tubes at 5 and 30min after cutting. **u**, Weighted means and SEWM (with variance weights) of cell volume after Rapamycin treatment on flat regions (dark blue squares) over time and on tubes (blue circles) over time after cutting. n≥16images/timepoint; N=3. **v**, Representative orthogonal views of Rapamycin-treated cells on flat regions or tubes at 5 and 30min after cutting. Scale bars=20 μm. N is the number of independent replicates.

To further test this hypothesis, we investigated membrane tension response to an epithelium rolling. Fluorescence lifetime instantaneously decreased upon rolling, earlier than the cell volume peak, and recovered to the initial value in about 10min (Fig. 4k,l), at the time when volume recovery was starting. Thus, tissue rolling correlated with a drop in membrane tension, followed by a volume increase that recovered once tension was back to its initial value.

As volume changes of epithelial cells were mostly accommodated on the apical side, we wondered if the apical actin belt was maintained during cell shape change. Moreover, actin is a general regulator of membrane tension^54^, and its modulation could explain the rapid drop in tension and its recovery that we observed in cells upon rolling. Interestingly, the apical/basolateral actin fluorescence ratio quickly decreased after rolling (Fig. 4m), due to a large decrease of apical actin staining, followed by a 20min recovery (Fig. 4n). We next wondered if actin depolymerization on the apical side was necessary to induce cell volume increase. When we used Jasplakinolide treatment to inhibit actin filament disassembly^55^, cell volume slowly and partially increased before recovery (Fig. 4o,p, Supplementary Fig. S4). Thus, inhibition of actin belt depolymerization counteracted cell volume increase. Most probably, the actin belt is involved in cell shape and membrane tension maintenance. Conversely, when treated with Blebbistatin, which disrupts myosin contractility, the dynamics of volume increase were similar to the non-treated condition (Fig. 4q,r, Supplementary Fig. S4). However, the resulting volume change was of ~20% (instead of ~40%), probably because cells treated with Blebbistatin had a higher initial volume. We then concluded that cell contractility had a minor role in the cell response to curvature change.

As actin, membrane tension and osmotic response are critically dependent on the mTORC1 & 2, we investigated their role in the response to curvature induction. We used Torin1 and Rapamycin treatments to respectively inhibit mTORC1&2^56^ and mTORC1^57^. In Torin1-treated cells, we observed a reduced volume increase (Fig. 4s,t, Supplementary Fig. S4) as compared to Rapamycin-treated cells (Fig. 4u,v, Supplementary Fig. S4). However, in both cases the volume peak was lower than in non-treated cells, and the recovery was delayed, suggesting that both cell volume increase and recovery involved both mTORC1 and mTORC2 signaling.

Altogether, our results show that a flat epithelium curled into a tubular shape actively and instantaneously responds by depolymerizing apical actin and decreasing plasma membrane tension (Fig. 5). This initial response is followed by a cell volume increase of ~40% 5min after rolling, and results from a swelling of the apical pole. Our data do not indicate whether actin depolymerization is the cause or the consequence of the membrane tension drop. However, the tension recovers while volume peaks, supporting that tension is re-equilibrated by cell swelling. Moreover, because volume increase happens a few minutes after tension decrease, and depends on mTORCs and ionic channels, we concluded that the volume increase is an osmotic swelling to restore membrane tension, in reaction to a stimulus perceived as a hypertonic shock. Then, cell volume returns to the initial values in about 20min. This process, associated to a recovery of the actin polymerization, is delayed when ion channels and mTORC1&2 activities are perturbed. This establishes the mechanical response to anisotropic curvature-induction as a regulated osmotic swelling.

**Figure 5.**
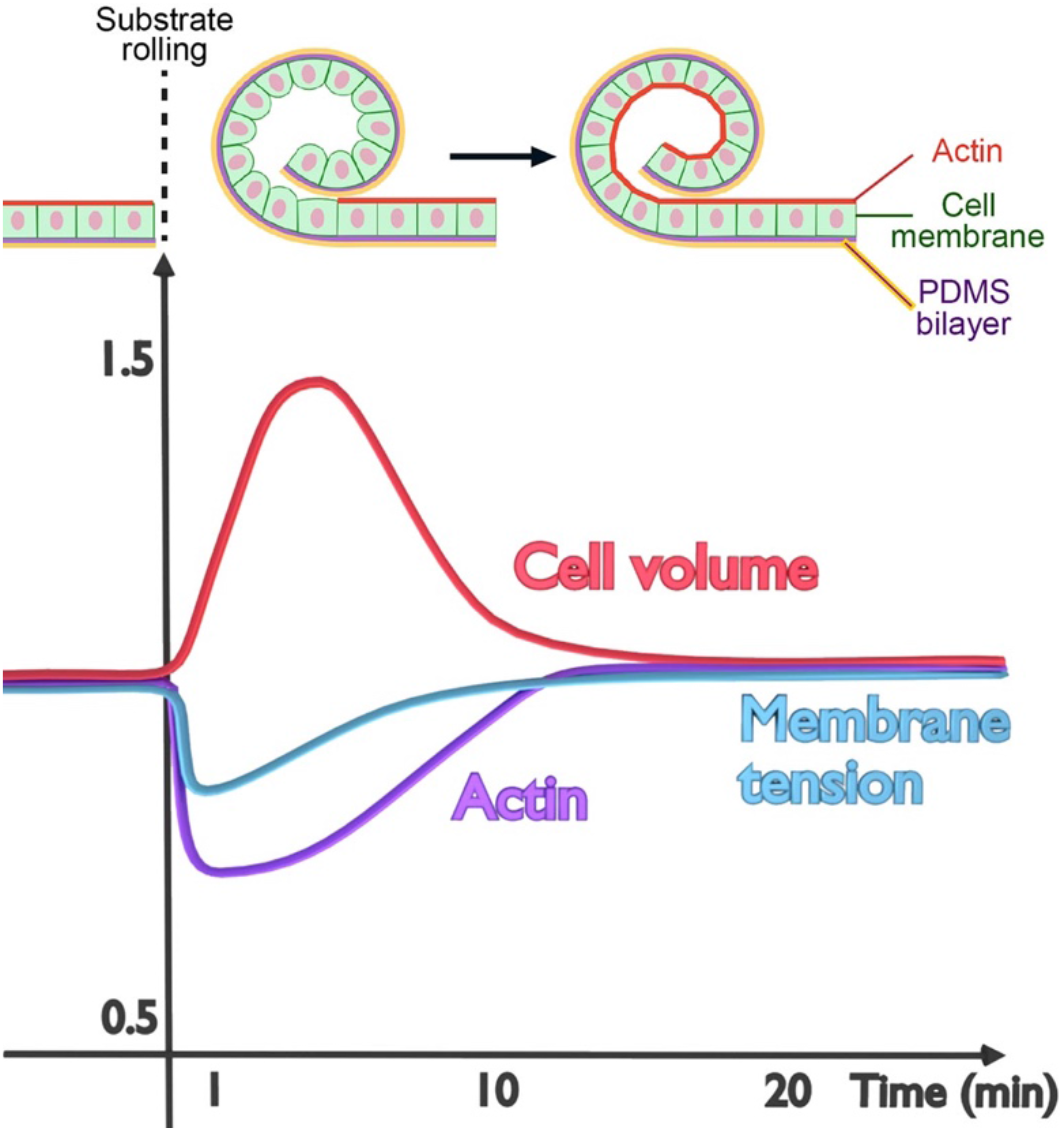
Mechanism controlling the transient cell volume increase upon rolling. Curvature generation of a MDCK cell monolayer induces both a membrane tension decrease and an actin depolarization on the apical side, which leads to a cell volume increase in a few-minute time range. The consequent membrane tension increase and actin reorganization allow recovery of the initial volume within 20min.

Finally, volume only fully recovers when apical actin is fully repolymerized. As volume recovery depends also on actin, our data support that the volume recovery is partially driven by actin polymerization. Interestingly, this response was only seen in large scale curvature induction, over curved distances of hundreds of microns, supporting the notion that only a collective reaction of cells to curvature can trigger this volume increase at the level of each single cell.

Overall, we show that epithelium is a unique active material that transiently swells upon rolling. Indeed, usual materials either reduce their volume or keep it upon bending or other curvature inducing deformations. These results may also bring new insights to the mechanisms of adaptation of very dynamics epithelia, such as lung alveoli, whose curvature regularly changes during the breathing cycle in a timescale of few seconds, acini or blood vessels. Speculating on the reason for such effect, we postulate that it participates in the necessary adaptation of cells to the new curvature: depolymerization of apical actin may be required to adapt the shape and the forces in the actin belt of each cell. The resulting drop in tension would cause the cell to swell, as observed in mitotic cells^58^, inducing the repolymerization of actin to recover volume^59^. Finally, the reformation of the actin belt will be adapted to the new curvature, recovering the force field that maintains epithelial cell cohesiveness. In this scenario, the active cell swelling is required so that cells do not lose volume and change shape too dramatically while the actin belt is disassembled. Thus, tissue remodeling requires a complex interplay between ion transportation, actin organization and volume regulation – possibly orchestrated by TORCs signaling^60^ – to adapt single cell mechanics to the new shape of the tissue.

## Methods

### Curvature-inducible substrate preparation

#### Self-rolling substrates with concave curvature

Ethanol-washed glass microscope slides (VWR, Pennsylvania, USA, Cat. No. 631-1550, 1mm thick) were diamond-cut to a size of 15×25 mm. Then, ~150 μl of Polydimethylsiloxane (PDMS) Sylgard 184 silicone elastomer (Dow Corning, Midland, USA, 10 wt% of curing agent) was poured on each piece of glass slide, vacuum-degassed for 10min, and cured at 100°C for 15min. This PDMS layer serves to obtain a sharp cut of material above it, allowing the blade to lean on it. The PDMS surface was then activated using a plasma cleaner (Harrick Plasma, New York, Cat. No. PDC-32G) for 45 s, though a 0.5 cm high PDMS mask with a central hole of 0.5 cm by 2 cm. ~200 μl (Sigma Aldrich, St. Louis, USA, Cat. No. G7041) of a 3% wt/vol of fish gelatin water solution were incubated >4h on the activated surface. Then, ~200μl of a solution of PDMS Sylgard 184, curing agent and Toluene (Fisher Scientific, Massachusetts, USA, Cat. No. T/2300/15) mixed in a 10:1:10 proportion were spin-coated onto the gelatin/PDMS surface (600 rpm/s, 3000 rpm, 50s), and reticulated on a hotplate at 110 °C for 30min. Then, ~200 μl of a second solution of PDMS Sylgard 184, curing agent, silicon oil (VWR, Pennsylvania, USA, Cat. No. 47 V 350) and Toluene mixed in a 5:5:0.5:10 proportion were spin-coated (600 rpm/s and, 1500 rpm, 50s), and reticulated as before. The substrate was finally incubated in isopropanol (Fisher Scientific, Massachusetts, USA, Cat. No. P/7500/15) overnight to extract the silicon oil. Then, on the day of cell seeding, the central part of the surface was plasma activated for 45s, through a 0.5cm thick PDMS mask with a central window of 0.5×2cm. Then, the activated surface was incubated with fibronectin (Life Technologies, Carlsbad, USA, Cat. No. 33016015, 1.8 μg cm^−2^ in phosphate-buffered saline (or PBS, Thermo Fisher Scientific, Waltham, USA, Cat. No. 18912014), 1h) to allow for cell adhesion.

#### Self-rolling substrates with convex curvature

The fabrication steps were the same than above, but the order of the two last layers was inverted, with spin-coating of the silicon oil-free PDMS solution being the last step.

#### Wavy substrates

As previously described^41^, silicone elastomer films (Silex Ltd, Linford, UK, super clear sheet, 0.5mm thick, #HT6240 40SH) were prestressed up to 100% of its length with homemade stretchers (composition: stainless steel and Teflon) and exposed with a laser beam (Gravotech, Rillieux-la-Pape, France, LS100 laser). The same stretchers were used for the experiments of compression and stretching of a flat cell monolayer.

### Cell culture and sample preparation

Madin Darby canine kidney II (MDCK-II) (ECACC, Cat. No. 00062107) and MDCK Myr-Palm-GFP H2B-mCherry cells were cultured in DMEM (Thermo Fisher Scientific, Waltham, USA, Cat. No. 61965026) supplemented with 10% fetal bovine serum (FBS, Thermo Fisher Scientific, Waltham, USA, Cat. No. 10270106, Lot No. 41G1840K), 1% Penicillin-Streptomycin (PS, Thermo Fisher Scientific, Waltham, USA, Cat. No. 15140122) and 1% nonessential amino acids (NEAA, Thermo Fisher Scientific, Waltham, USA, Cat. No. 11140035), at 37 °C and 5% CO_2_. MDCK Myr-Palm-GFP H2B-mCherry was generated as previously described^41^. Cell lines were regularly tested negative for contamination with mycoplasma (Eurofins GATC Biotech, Germany).

After trypsinization, cells were seeded onto the fibronectin-coated of the substrate at 0.6 10^6^ cells cm^−2^ (or 0.3 10^6^ cells cm^−2^ when indicated) and let for adhesion in the incubator for ~4h. Then, the medium was gently removed and samples were submerged of fresh warm medium. After ~20h, non-adherent cells were gently flushed away using a 1ml pipette tip. Samples were further processed as following.

#### Cells on self-rolling substrates

by cutting the PDMS bilayer with a scalpel in the central region, the gelatin swells and dissolves upon contact with the warm cell culture medium, releasing the pre-constraint and allowing for PDMS rolling. Thus, 2 symmetric tubes were produced per sample, kept in the incubator for defined timepoints after tube formation, and then fixed with 4% formaldehyde for 20min (PFA, Sigma-Aldrich, St. Louis, USA, Cat. No. F8775), washed three times with PBS and kept in PBS at 4 °C until imaging. In experiments where cell fixation was performed before cutting the substrate, samples were observed under the microscope at different timepoints after roll formation.

#### Cells on wavy substrates

After cells have been seeded and adhered until confluence to the prestressed and laser-treated elastomer films, the stretchers were brought back to a not-stressed position in order to produce the wrinkles in a few seconds and fixed in PFA at different timepoints as for the self-rolling substrates. For experiments where cells were compressed on flat elastomer films, cells were seeded on films pre-stretched by ~15%. The pre-stretch was rapidly released to induce compression, and cells were fixed in PFA at different timepoints as for the self-rolling substrates. For experiments where cells were stretched on flat elastomer films, cells were seeded on relaxed elastomer films mounted on stretchers, and rapidly stretched before fixation.

#### Hypo and hyper-osmotic treatments on flat substrates

Cells were observed at different timepoints after switching to a hypotonic (-~155mOsm) or hypertonic (+ ~510mOsm) solution. Hypotonic solutions were prepared by adding double distilled water in the cell medium (1:1). Hypertonic conditions were obtained by replacing the cell medium by a 0.5M sucrose (PanReac AppliChem ITW Reagents, Darmstadt, Germany Cat. No. A2211) solution. Live imaging was performed in Leibovitz medium without phenol red (Life Technologies, Carlsbad, USA, Cat. No. 21083027, supplemented with 10% FBS, 1% PS, 1% NEAA).

#### Hyper-osmotic treatments on self-rolling substrates

Cells were incubated in hypertonic (+~600mOsm, 0.5M sucrose) solution after cutting the substrate and fixed at different timepoints in PFA with the same concentration of sucrose than in the hypertonic cell medium.

#### Flat surfaces for side views of actin labelling

10×2×2mm PDMS parallelepipeds were glued by plasma activation on their largest side to the surface of glass-bottom dishes (MatTek, Bratislava, Slovakia, Cat. No. P35G-1.5-20-C). These samples were used to culture cell monolayer at the surface of PDMS, and easily image their side view as in tubes, allowing for comparable measurements of actin intensity.

### Drug treatments

Samples were incubated with drugs 30min before cutting the substrates to produce the tubes and incubated in the medium with the drugs until fixation, with the following concentrations: 100μM DCPIB (Tocris, Bristol, UK, Cat. No. 1540, 100mM stock solution in ethanol), 100μM EIPA (Tocris, Bristol, UK, Cat. No. 1540, 100mM stock solution in DMSO), 100μM Bumetanide (Sigma-Aldrich, St. Louis, USA, Cat. No. B3023, 100mM stock solution in DMSO), 100μM Furosemide (Sigma-Aldrich, St. Louis, USA, Cat. No. F4381, 100mM stock solution in DMSO), 200nM Jasplakinolide (Enzo Life Sciences, Lausen, Switzerland, Cat. No. ALX-350-275, 3mM stock solution in DMSO), 20μM Blebbistatin (Sigma-Aldrich, St. Louis, USA, Cat. No. B0560, 17 mM stock solution in DMSO), 250nM Torin1 (LC Laboratories, Woburn, USA, Cat. No. T-7887, 13mM stock solution in DMSO), 100nM Rapamycin (LC Laboratories, Woburn, USA, Cat. No. R-5000, 1mM stock solution in DMSO). DMSO (Dymethyl sulfoxide, Sigma-Aldrich, St. Louis, USA, Cat. No. D2650)

### Fluorescence labelling

After fixation, MDCK-II cells were stained with CellMask Deep Red Plasma Membrane Dye (1:1000; Molecular Probes, Thermo Fisher Scientific, Waltham, USA, Cat. No. C10046) and nuclei were labelled with Hoechst 33342 (1:2000; Thermo Fisher Scientific, Waltham, USA, Cat. No. 62249) for 15min at room temperature.

F-actin was labelled on live MDCK-II cells with 1.3μM SiR-Actin conjugated with Alexa Fluor 647 (Spirochrome, Stein am Rhein, Switzerland, Cat. No. SC001, 1mM stock solution in DMSO), incubated for 2 h. Cells were then fixed at different timepoints after cutting of the self-rolling substrates (see “Sample preparation”).

For immunostaining, after fixation, MDKC Myr-Palm-H2B cells were permeabilized using 0.1% saponin (Axonlab, Le Mont-sur-Lausanne, Switzerland, Cat. No. A4518,0100) in PBS and blocked in 1% wt/vol fish gelatin solution in PBS for 45min. Primary antibody anti-Ezrin (1:200, BD Biosciences, Allschwil, Switzerland, Cat. No. 610602) was incubated for 1h in 1% gelatin buffer. The secondary antibody donkey anti-mouse Alexa Fluor 488 (1:250; Thermo Fisher Scientific, Waltham, USA, Cat. No. A21202) was incubated for 45min in 1% gelatin buffer protected from light. Samples were washed three times with PBS after each antibody incubation. FlitptR probes were synthesized as previously reported^61^ or obtained commercially from spirochrome (Spirochrome, Stein am Rhein, Switzerland, Flipper-TR, Cat. No. SC020, stock solution 1mM in DMSO). 1h before imaging, MDCK-II cell medium was replaced with medium containing ~1μM of FliptR.

### Imaging

Fixed samples and PDMS tubes without cells (see “Image analysis”) were imaged in fluorescein dextran (Sigma, Aldrich, St. Louis, USA, Cat. No. FD40S, av. mol. wt. 40,000) solutions using an upright LSM 710 microscope with a water immersion Plan-Apochromat 20x/1.0 N.A. DIC objective (Carl Zeiss, Oberkochen, Germany), and operated with Zen 2012 software. Z-stack acquisitions were performed with Z-step of 0.5μm for cell imaging and of 1μm for imaging of PDMS tubes without cells. Confocal imaging of the samples labelled with actin or during osmotic shocks and Fluorescence Lifetime Imaging (FLIM) for lifetime measurements were performed with an inverted Nikon Eclipse Ti A1R microscope equipped with a time-correlated single-photon counting module from PicoQuant GmbH (Berlin, Germany). Excitation for FLIM was performed as described previously^53^ using a pulsed 485nm laser (PicoQuant, LDH-D-C-485) operating at 20MHz, and the emission signal was collected through a 600/50nm bandpass filter using a gated PMA hybrid 40 detector and a TimeHarp 260 PICO board (PicoQuant). A water immersion Apo LWD 40X/1.15 N.A. objective (Nikon) was used for imaging of actin-stained cells and FLIM acquisitions. A Plan Apo Lambda 100X/1.45 N.A. oil immersion objective (Nikon) was used for imaging of cells on flat substrates under hyperosmotic conditions for cell volume quantification. Z-stack acquisitions were performed with a Z-step of 0.6μm. During live imaging, cells were maintained at 37°C with 5% CO_2_ with a micro-incubator for thermal, CO_2_ and humidity control (Okolab, Pozzuoli NA, Italy). Confocal images of immunostained samples were obtained using Nikon Eclipse Ti microscope with an Andor Zyla sCMOS camera with 60X Plan Apo TIRF /1.49 N.A. DIC oil immersion objective (Nikon). The microscope was operated with Slidebook software. Z-stack acquisitions were performed with a Z-step of 0.3μm.

### Image analysis and segmentation

#### Cell volume quantification

Imaris 8.4.2 software (surface package) was used to segment fluorescent nuclei labelled with Hoechst or H2B-mCherry and calculate their centroid positions. To remove cells from the Z-stack edges that may have been partly cut, nuclei which position was equal or lower than 7μm from the image border were removed from the statistics. The centroid positions were then used as seeds for the segmentation of Z-Stacks of MDCK-II cells labelled with CellMask or Myr-Palm-GFP, and using LimeSeg^42^ plugin on FijiJ.

#### Actin intensity quantification

Icy software (point 2D tool) was used to select the regions of interest (ROI) of the apical and of the basolateral F-actin signal in order to quantify its intensity ratio.

#### Curvature quantification of PDMS tubes and wavy substrates

Radii of curvature were measured on self-rolling and wavy substrates at the PDMS surface where cells were supposed to adhere. On orthogonal views of the substrates, the chord of the arc (W) given by the PDMS substrate and the perpendicular bisector length (H) from the arc-chord were measured with the straight line tool in ImageJ. According to the chord theorem, the radius values (R) were then calculated as 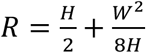

#### Fluorescence lifetime analysis

SymPhoTime 64 software (PicoQuant) was used to fit fluorescence decay data (from regions of interest, ROI) with a double exponential function, where the background was set to “0” value, and the intensity weighted average lifetime was extracted.

Independent replicates represent repeated experiments that include 3 samples per timepoint or condition and usually 1 or 2 images are acquired on each tube on the sample.

## Acknowledgements

Authors thank Nicolas Chiaruttini for his support for the data analysis performed with Limeseg, Tatiana Petitory, Adrien Leroy and Karine Anselme for their support during the first microfabrication experiments at the ISM, Giovanni Cappello and Pierre Nassoy for their critical reading of the manuscript. Authors also thank the Bioimaging Center of University of Geneva for their technical support and Roland Pellet and Maxime Domenjoz of the mechanical workshop of the University of Geneva for manufacturing the stretchers.

## Funding

A.R. acknowledges funding from the SystemsX EpiPhysX consortium, from the Swiss National Fund for Research Grants N°31003A_130520, N°31003A_149975 and N°31003A_173087, and the European Research Council Consolidator Grant N° 311536. V.L. acknowledges funding from the French National Research Agency ANR-17-CE18-0021-01 BioCaps.

## Author contributions

C.T. and A.R. designed the research. C.T. performed all experiments and image analyses. V.L. helped with the optimization of the rolling technique. L.B. and C.B-M helped with the theoretical models. C.T. and A.R. analyzed the data and wrote the manuscript, with editions from all co-authors.

## Competing financial interests

Authors declare no competing interests.

## Notes

### Competing Interest Statement

The authors have declared no competing interest.

