## Supplementary information for "Transient active osmotic swelling of epithelium upon curvature induction"

### Supplementary Figures

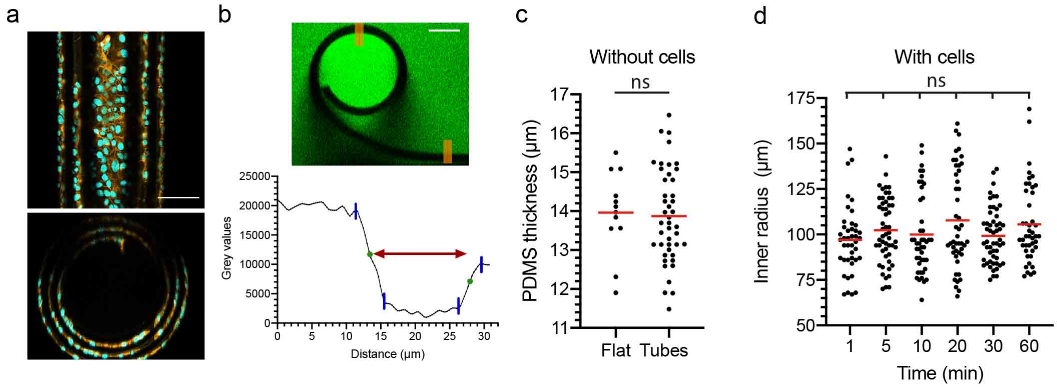


**Figure S1. a**, Example of formation of multi-rolls: top (top) and side (bottom) views of a 3 concentric PDMS tubes with cells, cell membrane (deep-red CellMask, orange), and nuclei (Hoechst, cyan). Scale bar = 100 µm. **b**, Method applied to measure the PDMS layer thickness of flat regions and of tubes. Top: PDMS layer identified by fluorescence exclusion (dextran solution, green). Scale bar = 50 µm. Bottom: example of plot profile got with a line thickness of 50 pixels (see the orange lines on the top image on the flat region and the tube). The PDMS width is measured between the 2 green circles, which represent the middle height between the min and the max peaks (blue lines). **c**, Distribution of PDMS thickness on flat regions and on tubes without cells. Mean values and SD: 13.96 ± 1.06 µm (Flat, n=12 images), 13.87 ± 1.20 µm (Tubes, n=44 images). N=3. **d**, Distribution of inner radii of tubes with cells as a function of time (minutes). Mean values and SD: 97.1 ± 18.4µm (1min, n=42 images), 102.3 ± 17.8 µm (5min, n=53images), 100.0 ± 21.8 µm (10 min, n=45 images), 107.8 ± 26.8 µm (20min, n=47images), 99.2 ± 15.2 µm (30min, n=53 images), 105.5 ± 21.5 µm (60min, n=44 images). N=3. N is the number of independent replicates, and the horizontal red lines stand for the mean values.

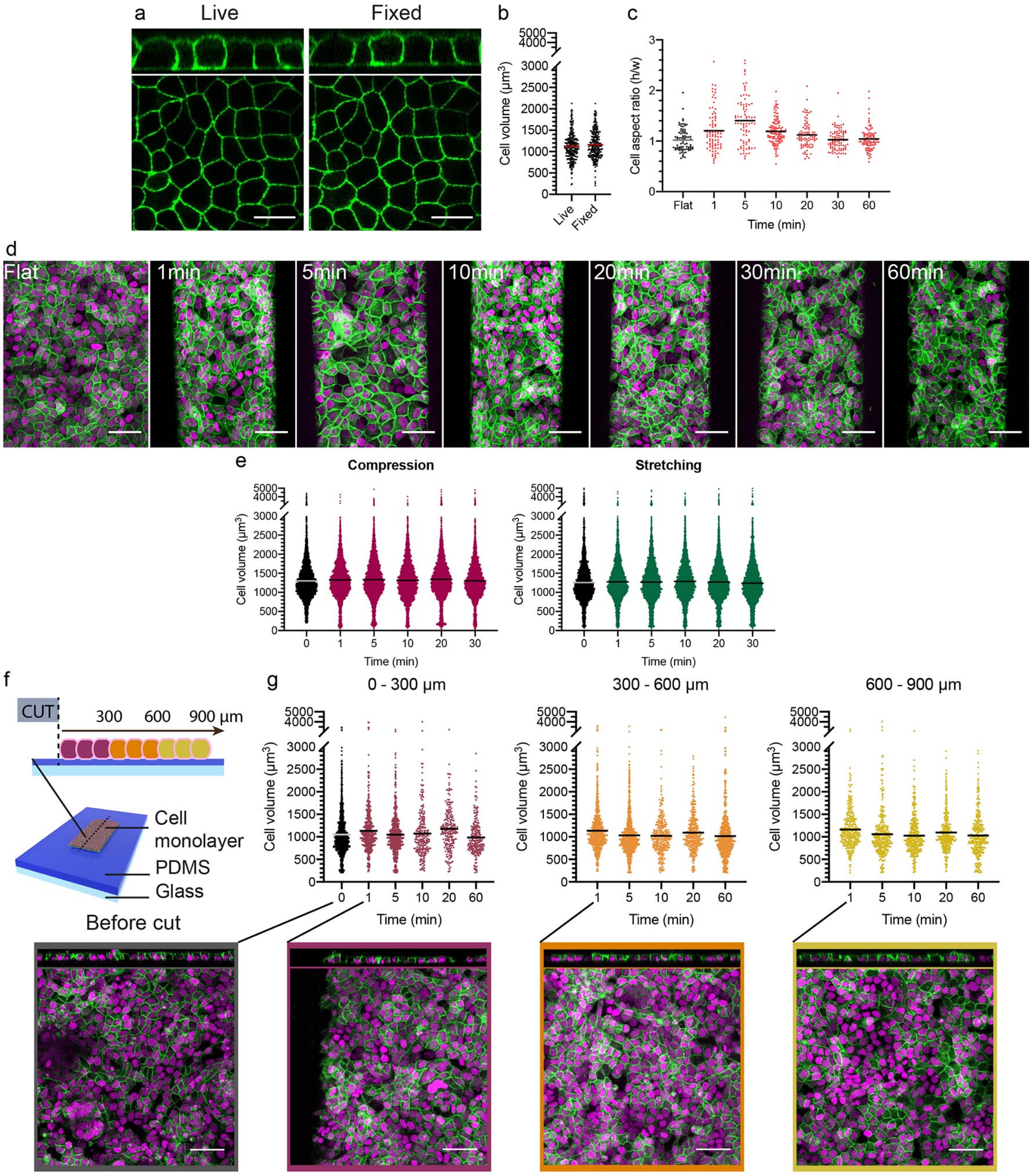


**Figure S2. a**, Representative orthogonal (top) and top (bottom) views of cell monolayers on a flat glass slide before ("Live") and after ("Fixed") chemical fixation, cell membrane (Myr-Palm-GFP, green). Scale bars=20µm. **b**, Distribution of cell volume on a flat glass slide before ("Live") and after ("Fixed") chemical fixation. n=250cells; N=2. **c**, Distribution of cell aspect ratio (height/width) on flat regions (black squares) and on tubes over time (minutes) after cutting (red squares). n≥81cells/timepoint; N≥3. **d**, From left to right: representative maximum-intensity z-projections of cell monolayers on a flat region and on tubes from 1 to 60 minutes after cutting, cell membrane (Myr-Palm-GFP, green), and nuclei (H2B-mCherry, magenta). Scale bars=50µm. **e**, Distribution of cell volume on an elastomer film before (black symbols) and after compression (left, magenta circles) or stretching (right, green circles) over time. n≥3406cells/timepoint; N≥3. **f**, Schematic of the regions observed over time from a cut of a cell monolayer on a flat PDMS substrate, where 0 µm indicates the position of the cutting. **g**, Distribution of cell volume on a flat PDMS substrate (without inner strain gradient) before (black symbols) and after cutting (from 0 to 300 µm from the cutting, magenta circles; from 300 to 600 µm from the cutting, orange circles; from 600 to 900 µm from the cutting, yellow circles) over time. n≥296cells/timepoint; N=1. Insets: representative orthogonal views (top) and maximum-intensity z-projections (bottom) of cell monolayers on a flat region before cut and at 0 to 300 µm, 300 to 600 µm and 600 to 900 µm from the cut at 1min after cutting, cell membrane (Myr-Palm-GFP, green), and nuclei (H2B-mCherry, magenta). Scale bars=50µm. N is the number of independent replicates, and the horizontal lines stand for the mean values.

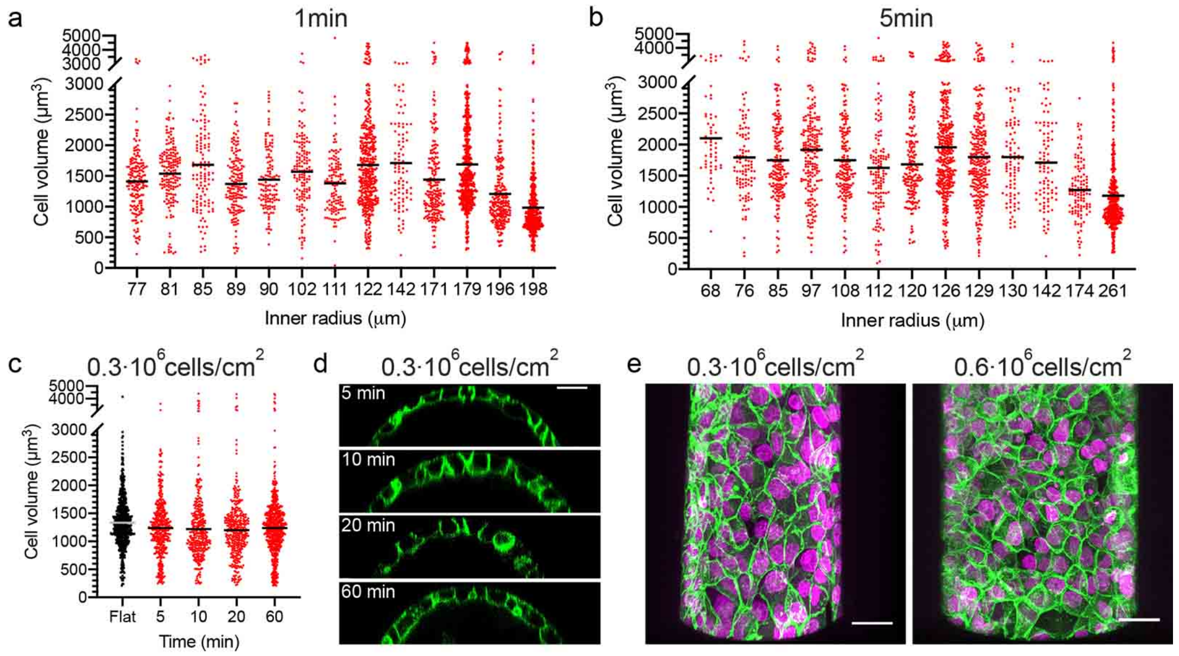


**Figure S3. a**, Distribution of cell volume on tubes 1min after cutting, as a function of the inner radius of the tube. n≥93cells/radius. **b**, Distribution of cell volume on tubes 5min after cutting, as a function of the inner radius of the tube. n≥58cells/radius. **c**, Distribution of cell volume on flat regions (black squares) and tubes (red circles) over time after cutting when the initial cell seeding density is of $0.3{\cdot}10^6$ cells/cm$^2$. n≥347cells/timepoint; N=2. **d**, Representative orthogonal views of cells on tubes at different timepoints after cutting, when the initial cell seeding density is of $0.3{\cdot}10^6$ cells/cm$^2$. Cell membrane (Myr-Palm-GFP, green). Scale bar=20µm. **e**, Representative maximum-intensity z-projections of cell monolayers on tubes 24 hours after cell seeding, with an initial cell seeding density of $0.3{\cdot}10^6$ cells/cm$^2$ (left) and $0.6{\cdot}10^6$ cells/cm$^2$ (right), cell membrane (Myr-Palm-GFP, green), and nuclei (H2B-mCherry, magenta). Scale bars=30µm. N is the number of independent replicates, and the horizontal lines stand for the mean values.

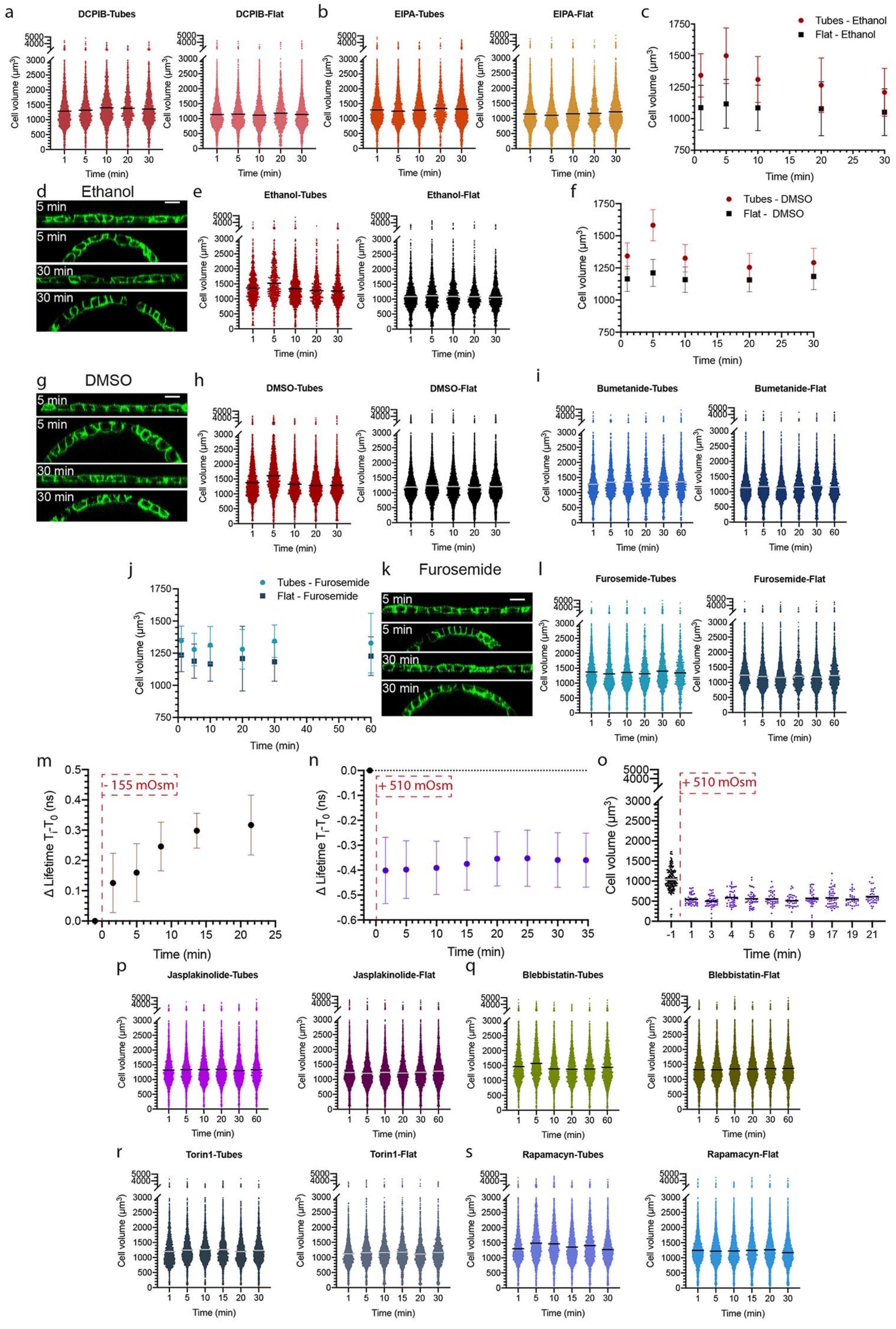


**Figure S4. a**, Distribution of cell volume after DCPIB treatment on tubes (left, dark pink circles) over time after cutting and on flat regions (right, light pink circles) over time. n≥1322cells/timepoint; N=3. **b**, Distribution of cell volume after EIPA treatment on tubes (left, light orange circles) over time after cutting and on flat regions (right, dark yellow circles) over time. n≥1317cells/timepoint; N=3. **c**, Weighted means and SEWM (with variance weights) of cell volume after ethanol treatment on flat regions (black squares) over time and on tubes (black circles) over time after cutting. n≥4images/timepoint; N=3. **d**, Representative orthogonal views of ethanol-treated cells on flat regions or tubes at 5 and 30 min after cutting, cell membrane (Myr-Palm-GFP, green). Scale Bar=20µm. **e**, Distribution of cell volume after ethanol treatment on tubes (left, red circles) over time after cutting and on flat regions (right, black circles) over time. n≥699cells/timepoint; N=3. **f**, Weighted means and SEWM (with variance weights) of cell volume after DMSO treatment on flat regions (black squares) over time and on tubes (black circles) over time after cutting. n≥17images/timepoint; N≥3. **g**, Representative orthogonal views of DMSO-treated cells on flat regions or tubes at 5 and 30min after cutting, cell membrane (Myr-Palm-GFP, green). Scale Bar=20 µm. **h**, Distribution of cell volume after DMSO treatment on tubes (left, red circles) over time after cutting and on flat regions (right, black circles) over time. n≥2360cells/timepoint; N≥3. **i**, Distribution of cell volume after Bumetanide treatment on tubes (left, blue circles) over time after cutting and on flat regions (right, dark blue circles) over time. n≥1311cells/timepoint; N=3. **j**, Weighted means and SEWM (with variance weights) of cell volume after Furosemide treatment on flat regions (blue squares) over time and on tubes (light blue circles) over time after cutting. n≥4images/timepoint; N=3. **k**, Representative orthogonal views of Furosemide-treated cells on flat regions or tubes at 5 and 30 min after cutting, cell membrane (Myr-Palm-GFP, green). Scale Bar=20µm. **l**, Distribution of cell volume after Furosemide treatment on tubes (left, light blue circles) over time after cutting and on flat regions (right, blue circles) over time. n≥912cells/timepoint; N=3. **m**, Mean values and SD of fluorescence lifetime (difference values of each timepoint – $T_i$ – with the initial timepoint – $T_0$) on flat regions before (-1min) and over time (minutes) after switching to a hypotonic solution (- ~ 155 mOsm). n≥2images/timpoint; N=1. **n**, Mean values and SD of fluorescence lifetime (difference values of each timepoint – $T_i$ – with the initial timepoint – $T_0$) on flat regions before (-1min) and over time (minutes) after switching to a hypertonic solution (+ ~ 510 mOsm). n≥6images/timepoint; N=1. **o**, Distribution of cell volume on flat substrates before (black circles) and over time (minutes) after switching to a hypertonic solution (+ ~ 510 mOsm, violet circles). n≥33cells/timepoint; N=1. **p**, Distribution of cell volume after Jasplakinolide treatment on tubes (left, light violet circles) over time after cutting and on flat regions (right, dark violet circles) over time. n≥1911cells/timepoint; N=3. **q**, Distribution of cell volume after Blebbistatin treatment on tubes (left, green circles) over time after cutting and on flat regions (right, dark green circles) over time. n≥2171cells/timepoint; N=3. **r**, Distribution of cell volume after Torin1 treatment on tubes (left, dark grey circles) over time after cutting and on flat regions (right, grey circles) over time. n≥1946cells/timepoint; N=3. **s**, Distribution of cell volume after Rapamycin treatment on tubes (left, dark blue circles) over time after cutting and on flat regions (right, blue circles) over time. n≥2199cells/timepoint; N=3. N is the number of independent replicates, and the horizontal lines stand for the mean values.

**Supplementary Information**

### Section 1: Estimation of the compressive force within the cell monolayer due to rolling of the PDMS substrate.

In this section, we use a continuum elastic description of the PDMS substrate and cell monolayer to estimate the pressure experienced by the cell monolayer due to the spontaneous curvature of the PDMS substrate, as well as the extent of the PDMS substrate unrolling due to the elastic resistance of the cell monolayer to bending.

We focus on the most curved region of the substrate + monolayer system in Fig. 1 of the main text, in which we assume its shape to be cylindrical. We also assume translational invariance along the axial direction of the cylinder, which allows us to focus only on the circular cross-section of the cylinder in the following. The substrate has a preferred radius of curvature $R_0 \simeq 100\ \mu m$ that is much larger than the thicknesses of the PDMS substrate and of the cell monolayer (both in the order of $\mathrm{h} \simeq 10\ \mu m$), therefore we describe substrate and cell monolayer as thin shells.

To estimate the pressure experienced by the cell monolayer due to the curvature of the substrate, we focus on a short time window (< 10 min) that follows the substrate's self-rolling, in which we assume that the cell monolayer behaves as a simple elastic material. When the cell monolayer sticks to the rolled substrate it is both bent and laterally compressed, and we postulate its surface energy density as

$$e_m = \frac{\mathrm{k_b}}{2}\left(\frac{1}{R}\right)^2 + \frac{\lambda}{2}\,\varepsilon^2 \tag{1}$$

where $k_b = 0.5\ \mu N \cdot \mu m$ (A. Trushko et al, Dev. Cell, 2020) is the monolayer bending rigidity, $R$ is its radius of curvature, $\lambda = 0.15\ \mu\mathrm{N}/\mu\mathrm{m}$ (A. Trushko et al, Dev. Cell, 2020) is its rigidity to lateral compression and $\varepsilon$ is its lateral strain. We estimate the strain as the relative variation of the length of the cell monolayer mid-line before ($l^0_{h/2}$) and after ($l_{h/2}$) bending, $\varepsilon = \left(l_{h/2} - l^0_{h/2}\right)/l^0_{h/2}$. We assume that the length of the cell-substrate contact line is the same before and after bending, and that $R$ is the radius of curvature there, in which case one finds that $\varepsilon = -h/(2R)$. Note that at $R = R_0$ we have $\varepsilon = 0.05$. We can now rewrite the surface energy density of the cell monolayer as

$$e_m = \frac{K_m}{2R^2}\ , \tag{2}$$

where $K_m = k_b + \lambda(h/2)^2$ is an effective bending rigidity of the cell monolayer. Note that $\lambda(h/2)^2 \simeq 3.8\ \mu N \cdot \mu m \gg k_b$, implying that most of the effective resistance of the cell monolayer to the rolling of the PDMS substrate comes from its resistance to lateral compression.

We now calculate the pressure required to stabilize the cell monolayer at a radius of curvature $R = R_0$, which is:

$$\mathrm{P}_{R=R_0} = \left(\frac{de_m}{dR}\right)_{R=R_0} = -\frac{K_m}{{R_0}^3} \simeq -4\ Pa\,. \tag{3}$$

In conclusion, we expect that shortly after self-rolling the substrate exerts a pressure in the order of $4\ Pa$ onto the cell monolayer in order to stabilize it.

Next, we estimate the extent of the PDMS substrate unrolling due to the elastic response of the cell monolayer. We start by evaluating the bending rigidity $K_s$ of the PDMS substrate, which can be written in terms of its thickness, Young modulus $E$ and Poisson ratio ν, by resorting to the classical theory of thin shells (Landau and Lifshitz, 1975, Theory of elasticity):

$$\mathrm{K}_s = \frac{E\, h^3}{12(1-\upsilon^2)}\,. \tag{4}$$

Using E = 1 MPa (Egunov e al, Soft Matter, 2016) and $\nu = 0.5$ for PDMS, we get $\mathrm{K}_s \simeq 100\ \mu N \cdot \mu m$. Then, we postulate the surface energy density of the substrate

$$e_s = \frac{\mathrm{K}_s}{2}\left(\frac{1}{R} - \frac{1}{R_0}\right)^2. \tag{5}$$

In the absence of a cell monolayer, the equilibrium radius of the substrate is $R_0$. In the presence of a cell monolayer, the equilibrium radius is the one that minimizes the total energy density $e = e_m + e_s$, which is

$$R^* = R_0 \frac{\mathrm{K}_s + \mathrm{K}_m}{\mathrm{K}_s} \simeq 104\ \mu\mathrm{m}\,. \tag{6}$$

To conclude, we expect that the unrolling of the substrate due to the resistance of the cell monolayer to bending deformations is $(R^* - R_0)/R_0 = 4\%$.

**Section 2: Estimation of the maximal thickening of the PDMS and cell layer upon rolling**

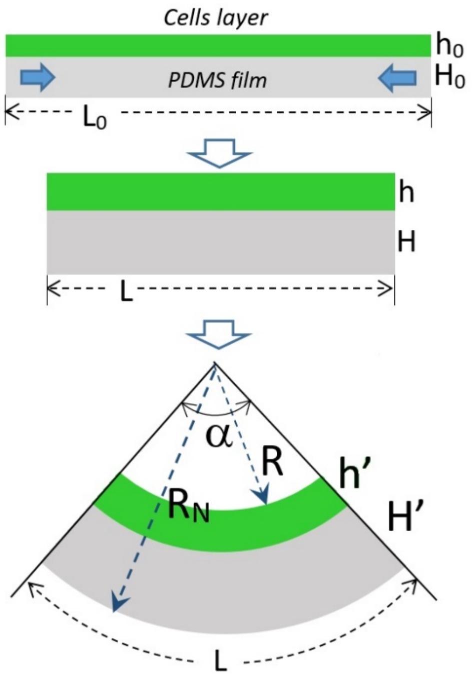


**Figure S5.** Geometric scheme for the estimation of the cell monolayer thickening.

To calculate the cells monolayer thickening due to geometrical reasons, and non-biological reasons, we split PDMS film relaxation on two steps: (i) in-plane shrinking of the film upon the release from the PDMS-glass substrate (Fig. 1a of the main text), and (ii) curling of the film due to internal bending moment in the PDMS film. We assume that the cell monolayer is much softer than the PDMS film, therefore we neglect its elastic response and assume it adapts its shape to the spontaneous curvature of the film. We model the substrate and cells monolayer as three-dimensional materials, with initial thicknesses $H_0$ and $h_0$, respectively, and long sides with typical initial lengths $L_0$ (Fig. S5). We determined experimentally that the PDMS layer experiences isotropic shrinking along its long sides when released from the substrate, when the rolling is hindered by placing a slippery glass slide atop of the film, such that the typical length of its long sides gets resized to $L = L_0(1 - \delta L)$, with $\delta L \simeq 0.02$. Let h and H be the thicknesses of the cell monolayer and PDMS, respectively, after lateral isotropic shrinking. Assuming that both cells and PDMS are incompressible, h and H can be calculated by enforcing the conservation of their volumes, yielding:

$$\begin{aligned} h &= \lambda h_0, \\ H &= \lambda H_0, \end{aligned} \tag{7}$$

where

$$\lambda = \frac{1}{(1-\delta L)^2}. \tag{8}$$

Let $H'$ ($h'$) be the thickness of the PDMS film (layer of the cells) after bending (Fig. S5). In order to find *the upper limit* on the thickening of the layer of cells due to the film bending, we make the following assumptions: (1) the neutral surface upon curling corresponds to the outer surface of the tube; (2) the tube does not elongate along its axial direction upon bending due to the Poisson effect. Under these assumptions, we calculate the thickening of the PDMS substrate and cell monolayer upon bending by enforcing the conservation of the areas of their cross-sections.

Consider a segment of the layer of the length $L$, which curls in a segment with the opening angle $\alpha$, inner radius $R$, and outer radius $R_N$, corresponding to the neutral surface, $R_N = R + h' + H'$ (Fig. S5). Conservation of the areas of the cross-sections of the PDMS and cells layers implies:

$$\begin{cases} \frac{\alpha}{2}[(R + h' + H')^2 - (R + h')^2] = LH \\ \frac{\alpha}{2}[(R + h')^2 - R^2] = Lh \end{cases} \tag{9}$$

Substituting $\alpha = \frac{L}{R_N} = \frac{L}{R+h'+H'}$ in (9), we get

$$\begin{cases}(R+h'+H')^2-(R+h')^2=2H(R+h'+H')\\ (R+h')^2-R^2=2h(R+h'+H')\end{cases} \tag{10}$$

which can be recast as

$$\begin{cases}2R(H'-H)+2h'(H'-H)+H'(H'-2H)=0\\ 2R(h'-h)+h'(h'-2h)-2hH'=0\end{cases} \tag{11}$$

Note that the system (11) is solved for H' = H and h' = h to leading order when $R \to \infty$, which corresponds to the case of flat layers. Given that in the experiments R is much larger than all other length scales, we exploit the large R limit to solve (11) perturbatively, close to the flat solution. We define the perturbed thicknesses H' $= H(1+\delta H/R)$ and $h' = h(1+\delta h/R)$, where $\delta H$ and $\delta h$ are of the same order of H and h. We substitute these expressions in (11) and keep only the leading order terms in the limit $R \to \infty$, which yields:

$$\delta H = \frac{H}{2} \tag{12}$$

$$\delta h = \frac{1}{2}(h+2H) \tag{13}$$

Using the experimental values (Fig. 1d, S1c, 2b) $R \approx 100\mu m$, $H_0 \approx 14\mathrm{\mu}m$, $h_0 \approx 10\mu m$, and $\lambda \approx 1.04$, we finally get $\mathrm{H}' \sim 15.6\ \mathrm{\mu m}$ and $h' \approx 12.5\mu m$. Therefore, the thickness increase due to shrinking and curling of the PDMS substrate does not exceed 12% for PDMS and 25% for the cell monolayer. Note that if a cell thickness increases of 25%, conservation of area implies that its width decreases of 20%, in this model.

However, the estimated PDMS layer thickening is comparable with the standard deviation of the experimental measurements ($14 \pm 1\mathrm{\mu}m$) and the PDMS thickness that we observed (Fig. S1c) is not significantly different before and after rolling. This confirms the prediction for a maximal PDMS thickness change upon self-rolling, which results in the same range than the experimental fluctuations.

The estimated cell thickening is less than the experimentally observed, suggesting that cell monolayers do not behave as elastic incompressible materials.

### Section 3: Estimation of the maximal compression of the PDMS layer upon rolling

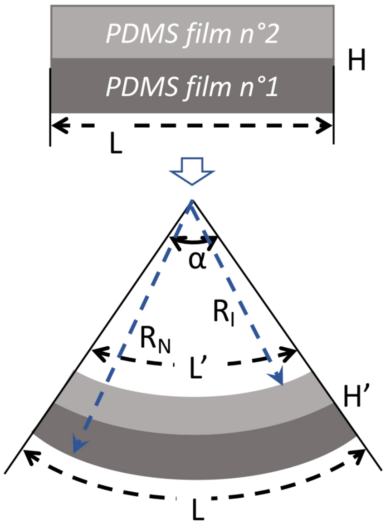


**Figure S6.** Geometric scheme for the estimation of the PDMS layer compression upon rolling.

To calculate the maximum PDMS layer compression due to the shrinking and the curling of the film, we assume that the neutral surface upon curling corresponds to the outer surface of the tube.

We consider a segment of the layer with initial length $L$ and thickness $H$, which curls into a segment with opening angle $\alpha$, thickness $H'$, inner radius $R_I$, and outer radius $R_N = R_I + H'$ (Fig. S6). We call L' the length of the inner face of the tube, such that, by construction:

$$\begin{cases} \alpha(R_I + H') = L \\ \alpha R_I = L' \end{cases} \quad (14)$$

We call $\Delta L = L - L'$, and obtain the relative variation of the PDMS layer length from system (14):

$$\frac{\Delta L}{L} = \frac{H'}{R_I + H'} \quad (15)$$

Substituting the estimated and the experimental values $H' \approx 15\mu m$ and $R_I \approx 100\mu m$ in (7) gives $\frac{\Delta L}{L} \approx 0.13$. Therefore, the compression of the PDMS layer due to shrinking ($\approx 2\%$) and curling of the PDMS substrate does not exceed 15%.

### Supplementary movies

**Movie 1. Curvature generation with a self-rolling substrate.** Bright-field real-time imaging showing a self-rolling substrate in cell medium with a confluent cell monolayer confined in the central region of the substrate. The cut with the scalpel induces spontaneous rolling of the PDMS bi-layer with the cells. Scale bar=0.5cm.

**Movie 2. Cell monolayer adaptation to curvature generation (top view).** Fluorescence confocal time-lapse of a rolled MDCK Myr-Palm-H2B cell monolayer on a tube after cutting of the substrate. Maximum-intensity z-projections overtime showing that cells are static in the time range of 60 min in which was observed the transient cell volume increase. Time format: h:min. Time interval = 1min until 25min, then 5min. Scale bar=100µm.

**Movie 3. Cell monolayer adaptation to curvature generation (side view).** Fluorescence confocal time-lapse of a rolled MDCK Myr-Palm-H2B cell monolayer on a tube after cutting of the substrate. Orthogonal view concatenation overtime showing that curvature generation does not induce an increase of extruded cells in the time range of 60 min in which was observed the transient cell volume increase. Time format: h:min. Time interval = 1min. Scale bar=50µm.
